# Emergent Entrainment and Predictive Dynamics in Bio-Inspired Spiking Neural Networks

**DOI:** 10.64898/2026.05.18.725874

**Authors:** Rodrigo Manriquez P, Sonja A. Kotz, Andrea Ravignani, Bart de Boer

## Abstract

Rhythm is a key building block of human music, speech and numerous other human activities. Understanding the computational substrates of rhythm perception requires models that bridge algorithmic function with biological implementation. We propose a physiologically grounded spiking neural network (SNN) framework to investigate the emergent representation and interpretation of auditory rhythms. Utilizing a recurrent SNN architecture trained on an auditory entrainment task, we characterize the network’s latent dynamics through the analysis of firing rates and membrane potential fluctuations. Our results demonstrate that simulated neural populations exhibit phase-locking to the stimulus beat, with endogenous oscillations driven by rhythmic input. We further show that anticipatory dynamics—characterized by pre-stimulus depolarization—emerge naturally from the network’s synaptic plasticity and temporal integration properties, rather than from explicitly defined oscillators. By treating network layers as functional analogs of cortical populations, this framework allows for the application of spectral and information-theoretic analyses typical of empirical electrophysiology. More in general, this approach establishes SNNs as robust exploratory tools for uncovering how predictive coding and rhythmic entrainment arise from the inherent constraints of biological neural computation.

## 1. Introduction

Rhythm lies at the core of human experience. The ability to synchronize with rhythmic patterns is a fundamental human capacity and enables essential social behaviors such as conversational turn-taking (Nguyen et al., 2020), coordinated dancing (Himberg et al., 2018) and musical performance in ensembles (Keller et al., 2014, Gordon et al., 2020). Moreover, rhythmic capabilities seem to not be exclusive to humans (Kotz et al., 2018; Ravignani et al., 2019), as rhythmic structure also plays a role in vocal interactions across several animal species. This highlights the broader biological significance of rhythm and motivates the study of how the brain encodes and processes rhythmic information. Despite its central role, the neural mechanisms underlying rhythmic processing are still not fully understood.

To better understand these mechanisms, a range of mathematical models and computational frameworks attempted to explain rhythmic perception and production (Large et al, 2023). Many of these approaches emphasize the predictability and periodic structure of rhythmic sequences. For example, frameworks based in Bayesian inference (Cannon, 2021) and predictive coding (Koelsch et al., 2019) propose that the brain constructs internal models that generate predictions about upcoming temporal events. Other models describe rhythm within a dynamical systems framework, like the Neural Resonance Theory (NRT) proposed by Large & Snyder (2009). NRT hypothesizes that empirically observable rhythmic features can be explained by neural level oscillations and that the coupling of neural oscillations gives rise to anticipatory behavior (Large et al., 2015). More broadly, and in alignment with the dynamical systems perspective, researchers have proposed neural-level descriptions of isochronous rhythms, defined as rhythmic sequences with evenly spaced temporal intervals. These approaches combine oscillator-based timing mechanisms with neural accumulators that integrate or count periodic activity to estimate temporal intervals and generate anticipatory responses (Bose et al., 2019; Egger et al., 2020).

Motivated by advances in artificial intelligence, additional attempts to explain observed neural dynamics have employed neural network (NN) models. Early studies of rhythm encoding relied on recurrent neural networks (RNNs) for beat modeling (Bock & Schedl, 2011), with subsequent work extending to more recent architectures such as transformers (Hung et al., 2022). The latter models achieve higher performance in beat tracking tasks at the cost of increased computational demands and reduced interpretability. In parallel, other approaches have explored neural networks not only as predictive tools but also as mechanistic models to the neural dynamics underlying rhythm processing. Further studies have employed biologically constrained RNNs to investigate firing rates in synchronization–continuation tasks (Zemlianova et al., 2024). The increased biological plausibility of this particular approach enabled direct comparisons with human tapping experiments (e.g., Gamez et al., 2019).

A logical next step toward a better representation of rhythm-related brain dynamics is to use Spiking Neural Networks (SNNs). SNNs offer increased biological plausibility by modeling neurons as discrete spiking units, more closely reflecting neural behavior and better align with established theories of neural entrainment and oscillations (Large & Snyder, 2009; Large et al., 2015). The main characteristic of SNN is that information is carried via precisely timed spikes (Maass, 1997), and although they typically lag behind traditional NNs in performance and are more challenging to train, they achieve comparable accuracy in tasks such as speech recognition (Yin et al., 2021; Bittar & Garner, 2022; Sun et al., 2023). Recent work further demonstrated that training SNNs on speech processing tasks can lead to the emergence of oscillatory neural dynamics and cross-frequency coupling across network layers (Bittar & Garner, 2024), supporting the use of spike-based architectures as mechanistic models for studying how neural oscillations emerge from network interactions.

We hypothesized that spiking neural networks naturally capture temporal dynamics and event timing, making them suitable models for exploring the neural mechanisms underlying rhythm processing. Utilizing a recurrent SNN architecture trained on an isochronous auditory rhythmic entrainment task, we characterized the network’s latent dynamics through the analysis of firing rates and membrane potential fluctuations. We investigated how rhythmic entrainment and anticipatory dynamics emerge in biologically grounded spiking neural networks trained on isochronous auditory input.

## 2. Methods

### 2.1 Spiking Neural Network

Here, we employed leaky integrate-and-fire (LIF) neurons, a computationally efficient extension of the original integrate-and-fire model (Lapicque, 1907) that enables gradient-based learning while retaining key features of spike-based computation (Neftci et al., 2019; Eshraigan et al., 2023). In a first-order LIF neuron model, the input is assumed to be a current injection *I*_*in*_[*t*] = *WX*_*in*_[*t*], where *X*_*in*_ [*t*] corresponds to the inputs from connections of the previous layer at timestep *t*, and *W* is a learnable weight matrix. Instead of passing the input through a nonlinear function, like in typical artificial NNs, the weighted inputs contribute to the membrane potential *U*[*t*], which decays over time at a rate β. When *U*[*t*] reaches a certain threshold *U*_*thr*_, the neuron fires a spike in the output *S*[*t*] and the threshold value is subtracted from *U*[*t*], resetting the neuron state. These dynamics can be summarized in the following equation:

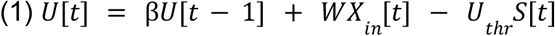

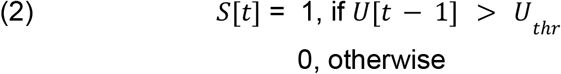

We also considered a version of the neurons, which included one-to-one recurrence, i.e., the output of each neuron is fed back as an input in the next time step. So, recurrence is implemented as follows:

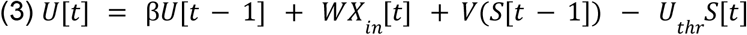

Where *V*(·) corresponds to a recurrent linear layer of the same size as *S*. In this LIF framework, *W*, β, and *U*_*thr*_ are learnable parameters. However, we only set *W* as learnable β fixing *U*_*thr*_, as learning the threshold has not been shown to yield any performance increase (Eshraigan et al 2023).

#### 2.1.1 Network architecture

We compared two spiking neural network architectures: a purely feedforward network (Eq. 1) and a recurrent network (Eq. 2), where recurrence was implemented independently for each neuron without shared recurrent connections within or across layers. This comparison allowed us to evaluate the effect of recurrence on network behavior.

Following a preliminary parameter exploration, both architectures consisted of three hidden layers containing 128, 256, and 128 neurons, respectively, consistent with previous work (Henkes et al., 2024). The output layer comprised a single leaky-integrator (LI) neuron with a sufficiently high threshold to prevent spike generation. Model output was obtained from the membrane potential of this neuron, following established approaches for regression tasks in SNNs (Henkes et al., 2024).

### 2.2 Auditory Input Pipeline

Auditory waveforms were converted into biologically motivated spike trains using a phenomenological model of auditory nerve transduction (Zilany et al., 2009, 2014). This model has been widely used in studies of hearing, speech processing, and rhythm perception (Carney, 2018; Zuk et al., 2018; Li et al., 2023). We simulated 32 high-spontaneous-rate fibers with characteristic frequencies distributed logarithmically between 125 Hz and 4 kHz. To increase the diversity of training samples, multiple spike-train realizations were generated for each auditory input while preserving the underlying firing statistics. Additional implementation details are provided in the Supplementary Material.

#### 2.2.1 Rhythmic Dataset

The dataset consisted of synthesized acoustic signals composed of trains of isochronous tonal pulses (Figure 1a). Pulses had a fundamental frequency ranging from 349 to 520 Hz in semitone steps, while different inter-onset intervals (IOIs) controlled the temporal spacing between consecutive events. Each pulse was shaped using an alpha-function-like envelope characterized by a decay-rate parameter (τ) that determined pulse duration (Figure 1c). During training, τ was randomized (Uniform distribution, τ = [0.04–0.08]) to increase variability, whereas a fixed value was used during evaluation.

**Figure 1.**
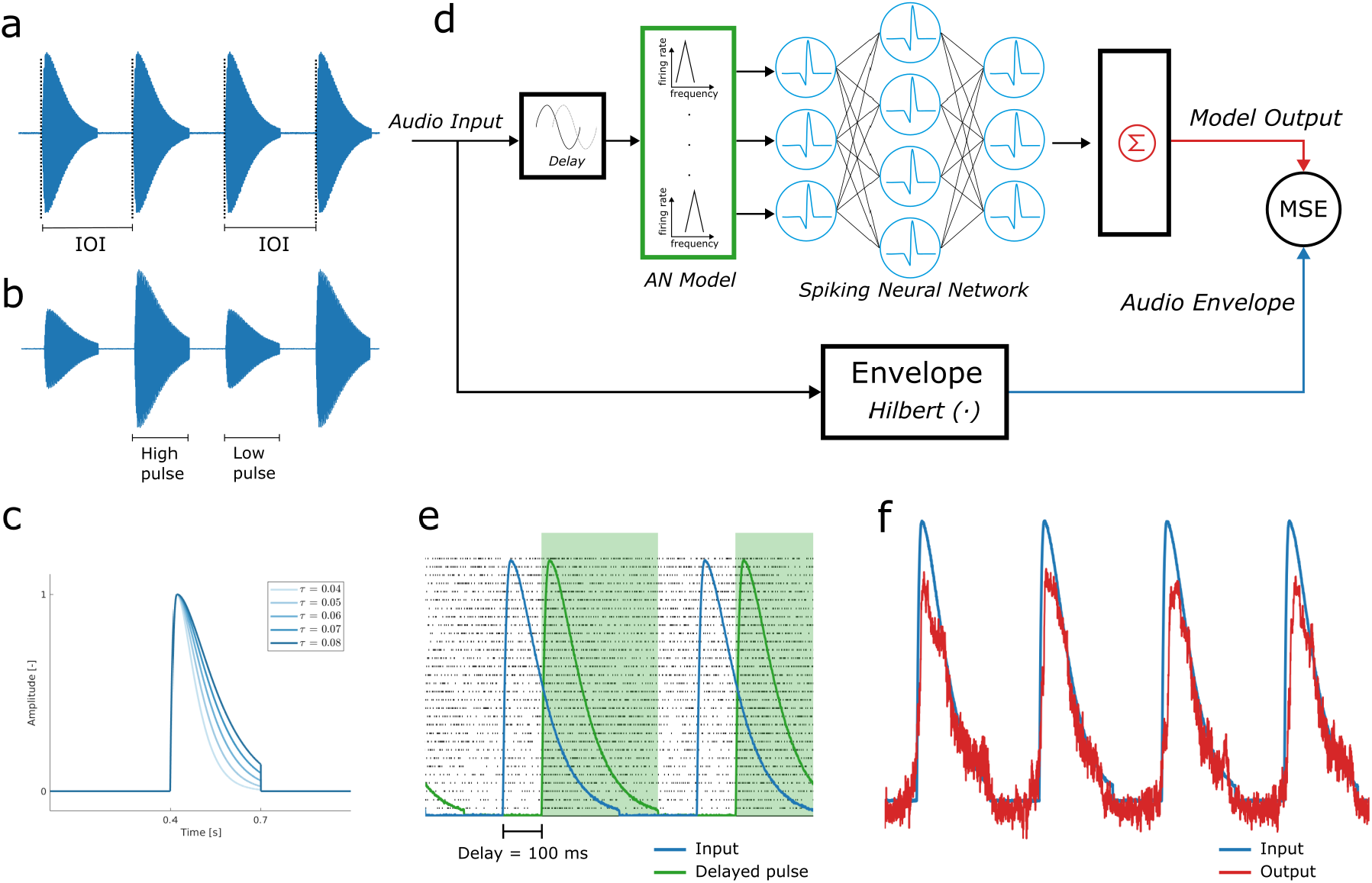
Proposed SNN framework. (a) Example of an isochronous pulse train used for training. (b) Example of the alternating-beats stimulus. (c) Pulse envelope with varying decay rates (τ). (d) Overview of the training pipeline: auditory signals are converted into spike trains, processed by the SNN, and trained to reconstruct the input envelope. (e) Illustration of the 100 ms input delay applied to spike trains. (f) Example network output compared with the target envelope.

Signals of 10 seconds duration were generated and converted into spike trains using the auditory nerve model. To increase the diversity of training samples, each signal was simulated 10 times, yielding multiple spike-train realizations. Training was performed on overlapping 3 s segments, while evaluation used the full 10 seconds signals.

Separate datasets were generated for IOIs of 0.4, 0.5, and 0.6 s, chosen to fall within the range of preferred human tempo (London, 2012). To evaluate responses to alternative rhythmic structures, we additionally generated an isochronous dataset with alternating high and low pulses, where high pulses had twice the amplitude and were two semitones higher in fundamental frequency than low pulses (Figure 1b). Finally, to assess generalization, an additional dataset was generated with IOIs ranging from 0.45 to 0.55 s in steps of 0.01 s.

#### 2.2.2 Delay

Signal transmission in biological neural systems is not instantaneous, introducing delays that accumulate across successive processing stages. Such delays have been proposed as a key factor underlying anticipatory behavior in rhythmic tasks, as delayed dynamical systems can exhibit anticipation of an external driver (Dubois, 2001; Large & Snyder, 2009). Experimental work further suggests that anticipation may emerge because of these delays rather than despite them (Stephen et al., 2008; Stepp & Turvey, 2010). To evaluate this effect, we introduced a fixed 100 ms delay to the input spike trains (Figure 1d), approximating the latency of early cortical auditory responses (Luck, 2014), which are sensitive to stimulus onset and temporal predictability (Schwartze et al., 2013).

### 2.3 Training and Validation

Computational implementation and training were performed in Python. For ease of neural network implementation, we used the SNNtorch library (Eshraigan et al., 2023), which enabled spiking neuron modeling and training using PyTorch. One advantage of SNNtorch is that it allows training spiking neural networks using backpropagation through time by implementing surrogate gradient descent (SGD). SGD overcomes the non-differentiable nature of spikes by replacing the true gradient with a smooth surrogate (Neftci et al., 2019). To further stabilize training, we employed Threshold-Dependent Batch Normalization (tdBN) (Zheng et al., 2021), a normalization method specifically designed for spiking neural networks. SGD has been successfully demonstrated to effectively train SNNs across different applications (Dampfhoffer and Mesquida, 2024; Sun et al., 2025), and, more importantly, it has recently been shown that SNNs trained with SGD can extract timing-base information and coding in speech tasks (Yu et al., 2025).

Networks were trained for 8 epochs, as preliminary testing showed that accuracy stabilized by this point. A batch size of 10 was used, together with the Adam optimizer and a learning rate of 10^−4^. The sampling frequency was set to 2 kHz, as higher frequencies substantially increase computational complexity and training time. Training was performed using a single NVIDIA GeForce RTX 3070 Ti GPU.

The output of the model corresponded to the estimated auditory envelope of the input. Model performance was quantified using the mean squared error (MSE) between the input envelope and the membrane potential of the integrating neuron in the output layer (Figure 1d). We hypothesized that this envelope should capture the main rhythmic features of the input.

### 2.4 Analysis of SNN dynamics

Understanding how artificial neural networks represent and transform information is challenging (Lipton, 2018). Neural network interpretability has emerged as its own research area, spanning multiple methodologies and facing difficulties that vary across architectures and tasks (Fan et al 2021). In particular, for SNNs there is no consistent way of analyzing their dynamics, as they are often evaluated in terms of their accuracy or through task-specific methods. Given that we wanted to exploit their potential as biological models of the brain, we borrowed techniques often used to analyze neural populations in real neurons, with the aim that analogous analyses can be performed.

#### 2.4.1 Layer Analysis

Given their hierarchical structure, we hypothesized that each layer in an SNN can be treated as an individual neural population. Previous research has shown that activity across layers of neural networks exhibits similarities to neural activity in the brain, particularly in the auditory cortex (Kell et al., 2018). For each neuron, we extracted both firing rates and membrane potentials over time. To characterize population-level dynamics while reducing dimensionality, we applied Principal Component Analysis (PCA) to the activity of each layer, following approaches commonly used to study neural population trajectories in sensorimotor timing tasks (Gamez et al., 2019). This allowed us to visualize low-dimensional representations of neural activity and investigate how population dynamics evolved in relation to the rhythmic input. Additional details regarding the PCA implementation are provided in the Supplementary Material.

#### 2.4.2 Mean Asynchrony

After training, we evaluated the ability of the SNN to synchronize with the input by comparing predicted and target onset times. To this end, we extracted onset timestamps using an energy-based onset detection algorithm and computed the mean asynchrony between the network output and the input signal. Mean asynchrony provides a measure of temporal alignment, with negative values indicating anticipatory responses and positive values indicating delayed responses relative to the reference onset.

## 3. Results

### 3.1 Isochrony

Figure 2 shows the network output for IOIs of 0.4, 0.5, and 0.6 seconds in both network types. The SNN output (orange) tracked the envelope of the acoustic input (blue). Accuracy (MSE) results are shown in Table 1. Overall, recurrent networks outperformed feedforward networks, although both architectures successfully entrained to the isochronous signals. The recurrent network more accurately reproduced the envelope shape, whereas the feedforward network showed less precise onset alignment and envelope tracking.

**Table 1.**
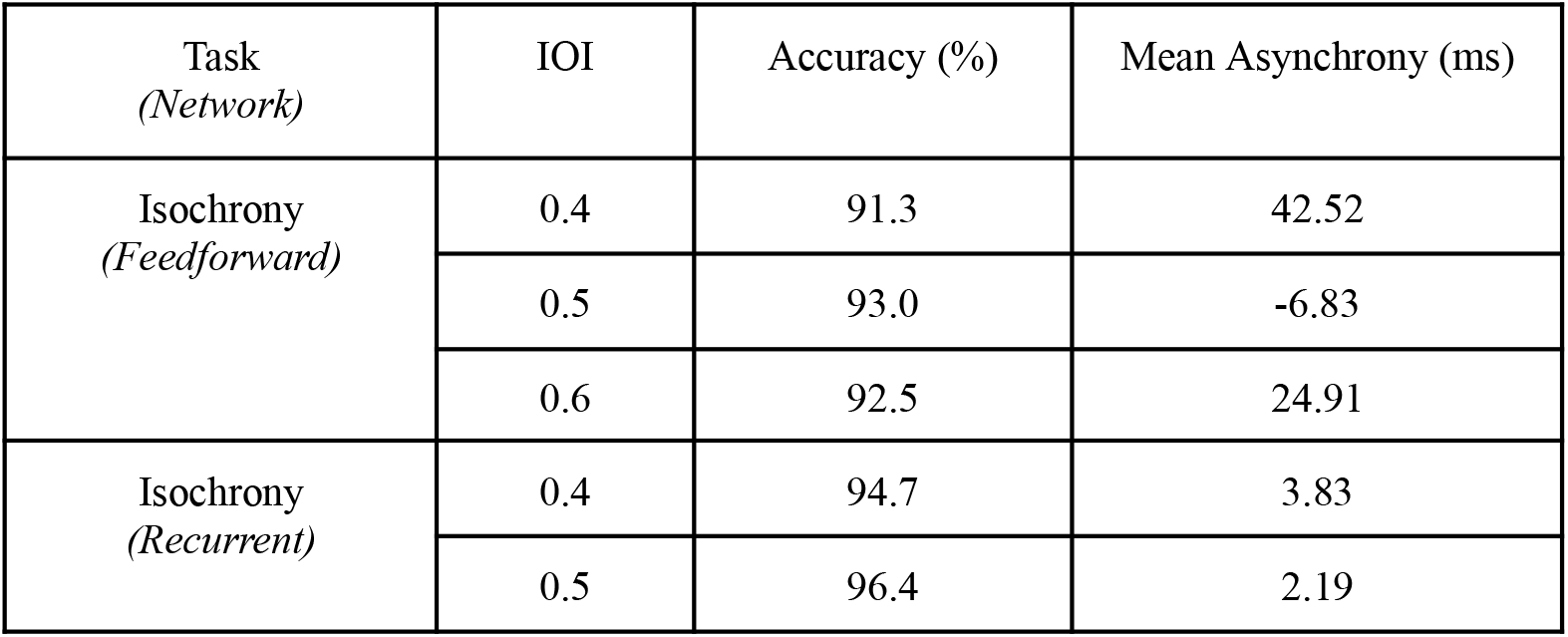
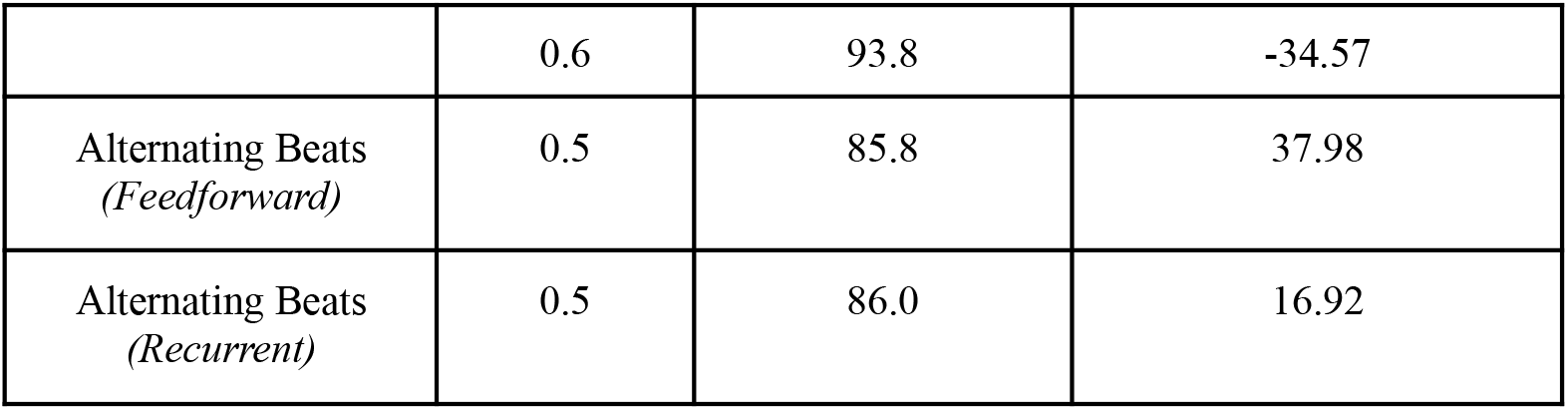
Accuracy and mean asynchrony for networks trained on isochronous tempi in different tasks. Overall, networks without recurrence underperformed compared against the ones with recurrence.

| Task<br>(Network) | IOI | Accuracy (%) | Mean Asynchrony (ms) |
| --- | --- | --- | --- |
| Isochrony<br>(Feedforward) | 0.4 | 91.3 | 42.52 |
|  | 0.5 | 93.0 | -6.83 |
|  | 0.6 | 92.5 | 24.91 |
| Isochrony<br>(Recurrent) | 0.4 | 94.7 | 3.83 |
|  | 0.5 | 96.4 | 2.19 |
|  | 0.6 | 93.8 | -34.57 |
| Alternating Beats<br>(Feedforward) | 0.5 | 85.8 | 37.98 |
| Alternating Beats<br>(Recurrent) | 0.5 | 86.0 | 16.92 |

**Figure 2.**
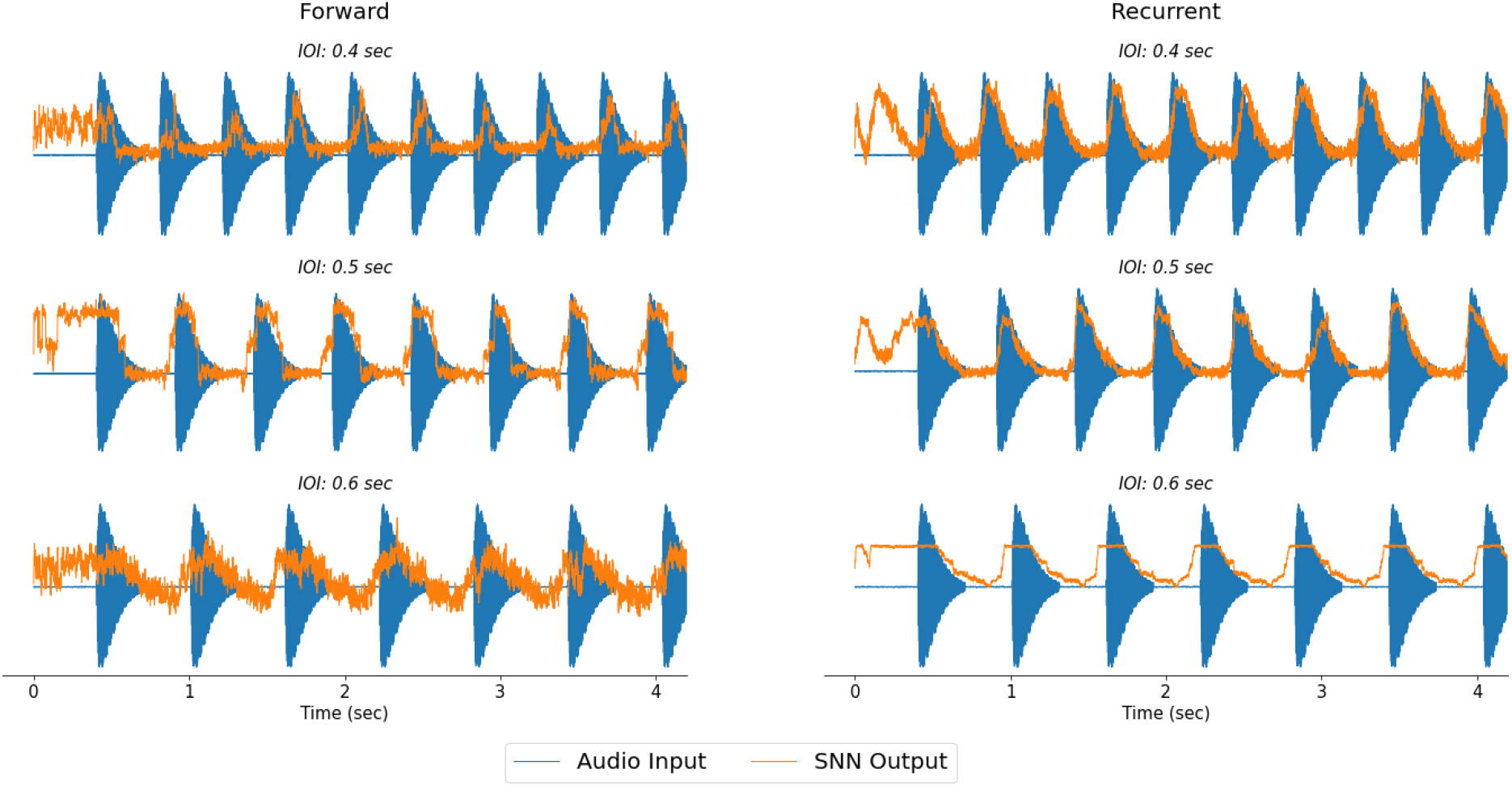
Results for Feedforward (left) and Recurrent (right) networks, trained on IOIs 0.4 (top), 0.5 (mid) and 0.6 (bottom).

To highlight the predictive nature of the SNN, Figure 3 shows a pulse omission, where one pulse in the isochronous sequence was removed. Both networks predicted the missing pulse but failed to sustain the oscillation, becoming “stuck” at a high output value until a subsequent pulse arrived and restored synchronization. When the stimulus stopped entirely, oscillatory activity disappeared and the output became noisy.

**Figure 3.**
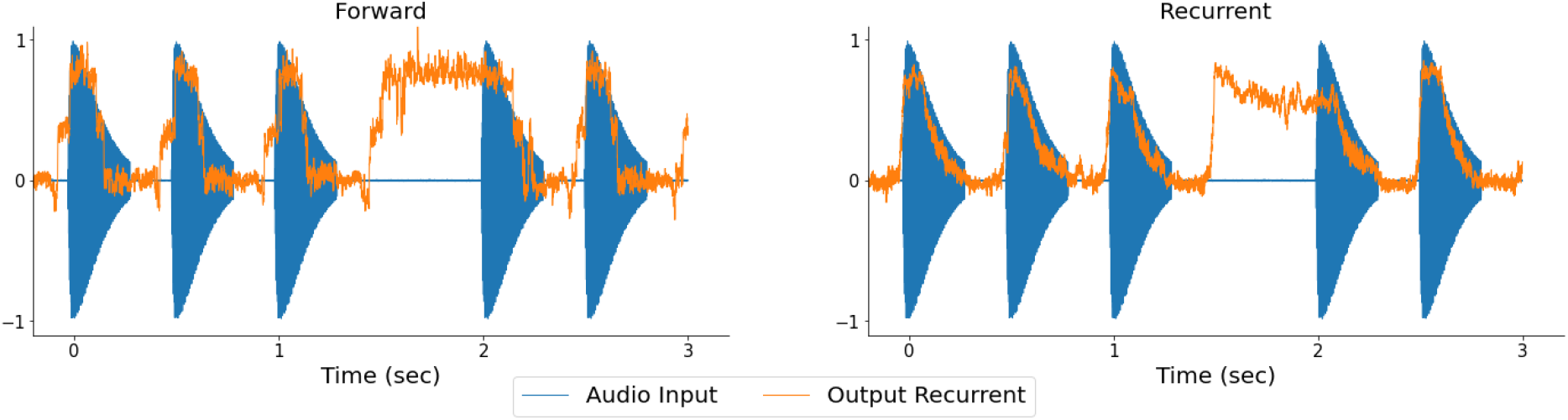
Results for feedforward (left) and recurrent (right) networks, trained on an isochronous dataset with IOI=0.5. Here, a beat is missing in the input, with the network predicting the missing onset.

The results for the alternating beat task are shown in Figure 4. In this case, both networks could entrain the output to the beat, aligning with the onsets appropriately, but were unable to match the amplitude accordingly.

**Figure 4.**
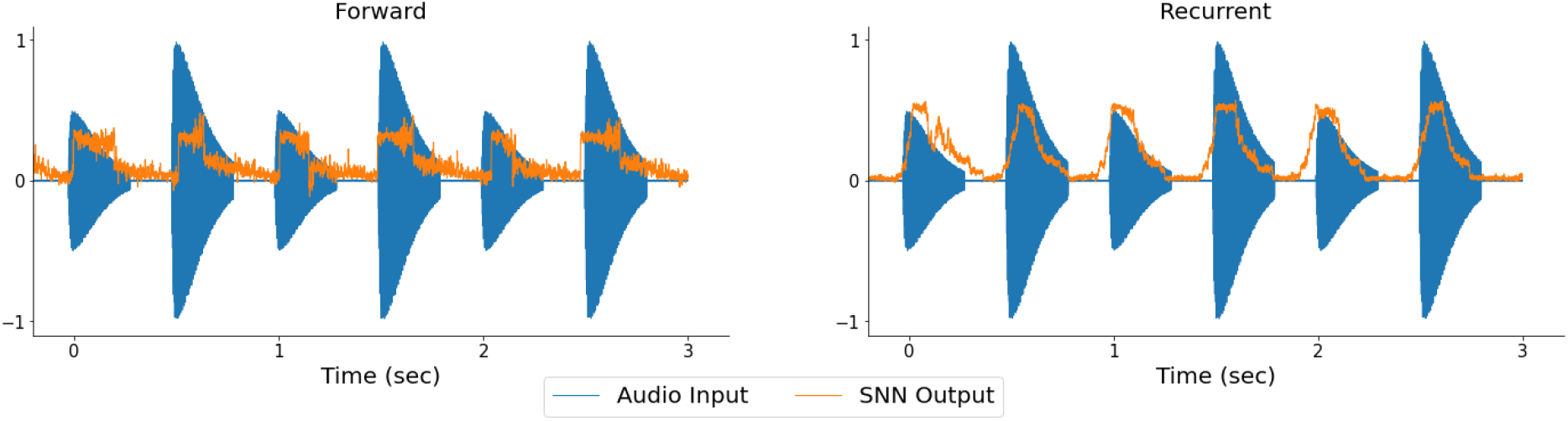
Results for feedforward (left) and recurrent (right) networks trained on IOIs with alternating beats. In both cases, the networks were able to predict the underlying beat, similarly to the isochronous condition, but failed to adjust the amplitude accordingly.

To further analyze network behavior, we examined firing rate and membrane potential dynamics across all layers. Figure 5 shows the firing rates of the recurrent SNN trained and evaluated on an IOI of 0.5 s over a 10 s interval. Neurons are displayed along the vertical axis and grouped by a clustering algorithm, with brighter colors indicating higher firing rates. Acoustic onsets are shown as vertical reference lines; due to the encoding delay, pulses reached the network 100 ms later.

**Figure 5.**
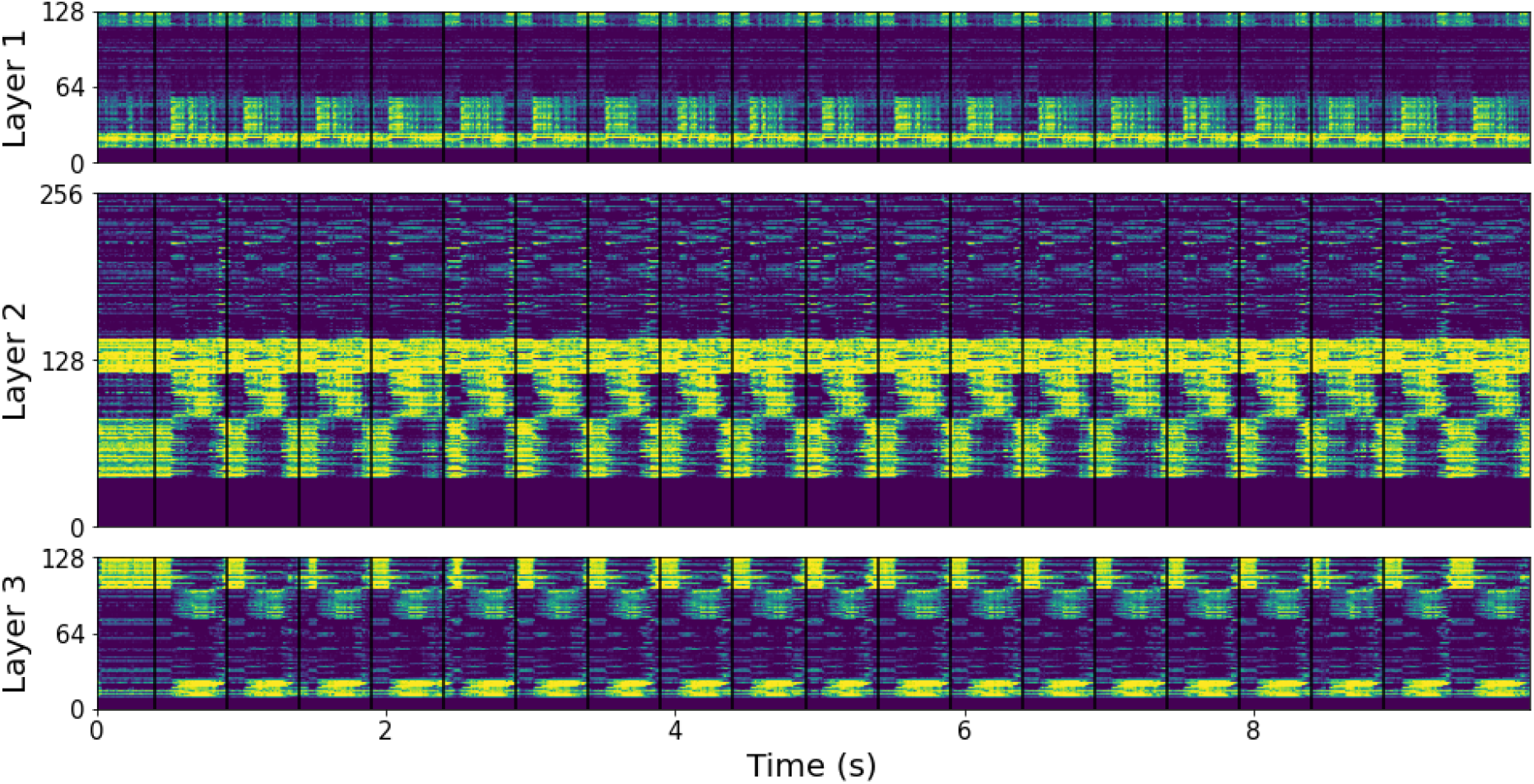
Firing rates of spiking neurons across network layers. Individual neurons are sorted along the vertical axis, and brighter colors indicate higher firing rates. Distinct alternating firing patterns were observed across clusters, while silent neurons (no firing activity) were clustered at the bottom.

Some neuronal clusters became active following pulse arrival, whereas others were active between pulses and silent at onset, producing alternating patterns of activity and silence. This organization was less apparent in clusters with low firing rates. Additionally, a population of silent neurons that never fired was observed across all layers and clustered at the bottom of the figure. Similar patterns were consistently observed in all networks trained on isochronous pulses.

Although the alternating firing patterns in Figure 5 suggested synchronization to the input, we further characterized population dynamics by computing principal components of the firing rates and membrane potentials. Figure 6a shows the first three principal components for Layer 3 of the same network. Vertical lines indicate predicted onsets, and dashed lines indicate the onset of the delayed spike trains. The components oscillated and entrained to the incoming beat. Moreover, firing-rate components remained largely in phase, whereas membrane-potential components exhibited phase differences. This is illustrated in the trajectory plots (Figure 6b), where membrane potentials (orange) formed circular trajectories, while firing rates (green) formed narrow ellipses, consistent with their smaller phase differences.

**Figure 6.**
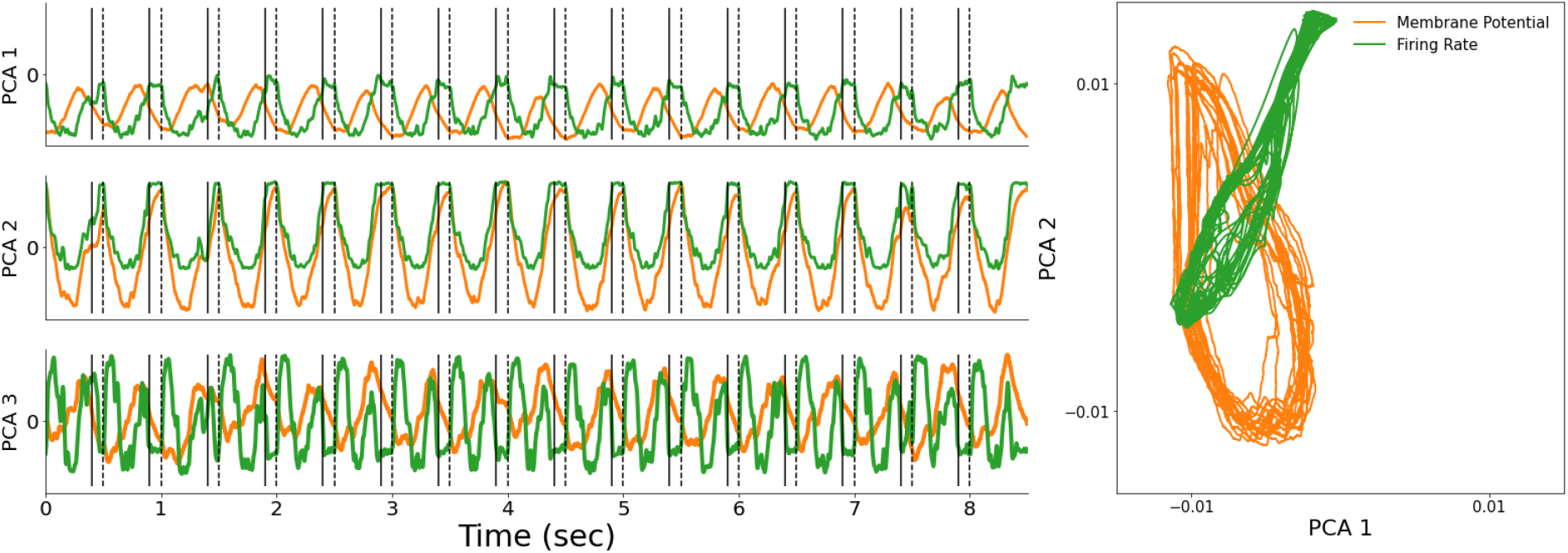
Principal component analysis of Layer 3 in the recurrent SNN for an input with an IOI of 0.5 s. Membrane potentials are shown in orange and firing rates in green. (a) First three principal components. Solid lines indicate pulse onsets and dashed lines the corresponding delayed spike trains. The first two components explained 84% of the variance in firing rates and 97% of the variance in membrane potentials. (b) PC1–PC2 trajectories.

Through these simulations, we showed that neural subpopulations entrained to the beat of the acoustic input. This entrainment was reflected in the oscillatory behaviour of the membrane potentials, which can be interpreted as an analogue of electrophysiological activity. Such behaviour is consistent with electrophysiological recordings in humans, where neural activity entrains to the beat frequency of musical rhythms (Nozaradan et al., 2016). Likewise, circular trajectories have been reported in auditory cortical populations of macaques during finger-tapping tasks (Gamez et al., 2019).

The task also required the SNN to anticipate upcoming onsets. Evidence for this mechanism can be observed in the second firing-rate principal component (Figure 6a, middle row), which increased and saturated near the predicted onset. To further characterize these dynamics, we computed the PSTH, averaging firing rates (green) and membrane potentials (orange) for the same example. Results are shown in Figure 7. Four neuronal clusters are shown, representing distinct subpopulations within each layer. Vertical dashed lines indicate the onset to be predicted, while the encoded pulse information arrived 100 ms later.

**Figure 7.**
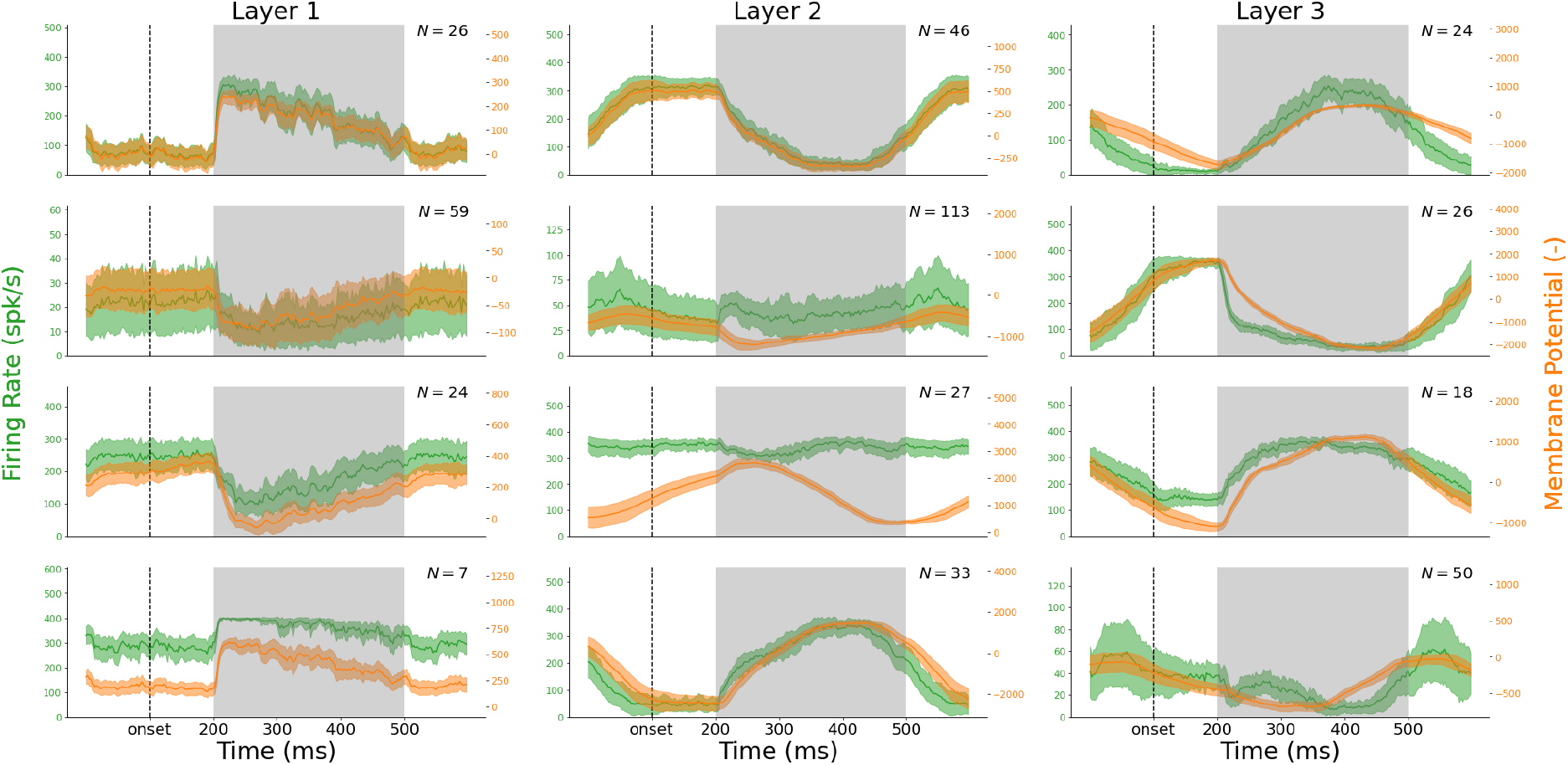
PSTHs of the mean firing rates (green) and membrane potentials (orange) of subpopulations of neurons in the third layer of the network with recurrence, obtained over 18 repetitions of a signal with IOI of 0.5 sec. Shaded curves depict 95% confidence interval (CI). Vertical dashed lines indicate the predicted onset (t = 100 ms), and a darker gray background indicates the presence of delayed pulse information (t = 200–500 ms).

We found that the average membrane potential of some neuronal subpopulations increased (or decreased) exponentially and saturated near the predicted onset before resetting in response to the delayed pulse. This suggests that predictive capabilities rely on the gradual accumulation of input spikes toward a threshold, enabling the network to anticipate upcoming onsets. These dynamics were observed across all trained networks, regardless of recurrence, differing mainly in waveform shape and polarity. However, they were not present in every layer; for example, in Figure 7 (left row), all neuronal clusters in the first layer responded only to the delayed pulse.

Pulse onsets predicted by the SNN emerged from the entrainment of neural populations across layers. Further testing showed that oscillatory activity disappeared when the rhythmic input was removed or replaced by noise (Supplementary Figure 1). This suggests that the network did not develop an internal oscillator tuned to the stimulus frequency; instead, anticipation and synchronization emerged in response to rhythmic input. Unlike beat-generation models that sustain rhythmic activity in the absence of input (Bose et al., 2019; Zemlianova et al., 2024), the network did not exhibit autonomous oscillations after stimulus offset.

### 3.2. Generalization

So far, we have shown that SNNs can entrain to the IOI of an acoustic input. We then evaluated the ability to synchronize across multiple IOIs and adapt to different inputs . Figure 8 shows the results of a generalization task for both types of networks, with the same network being evaluated across different IOIs. Accuracy results for the different IOIs are shown in Table 2. Similar to the previous task, both networks could entrain to the input rhythm, with the recurrent case again performing better overall. Importantly, a single SNN could synchronize to multiple rhythms.

**Table 2.** Accuracy and mean asynchrony for networks trained on the generalization task. Performance of the network with recurrence was better than without recurrence.

| Task<br>(*Network*) | IOI | MSE (%) | Mean Asynchrony |
| --- | --- | --- | --- |
| Generalization<br>(*Feedforward*) | 0.45 | 94.1 | 14.11 |
|  | 0.5 | 93.6 | -21.8 |
|  | 0.55 | 92.5 | -46.9 |
| Generalization<br>(*Recurrent*) | 0.45 | 94.4 | 25.11 |
|  | 0.5 | 96.0 | -14.45 |
|  | 0.55 | 95.0 | -45.09 |

**Figure 8.**
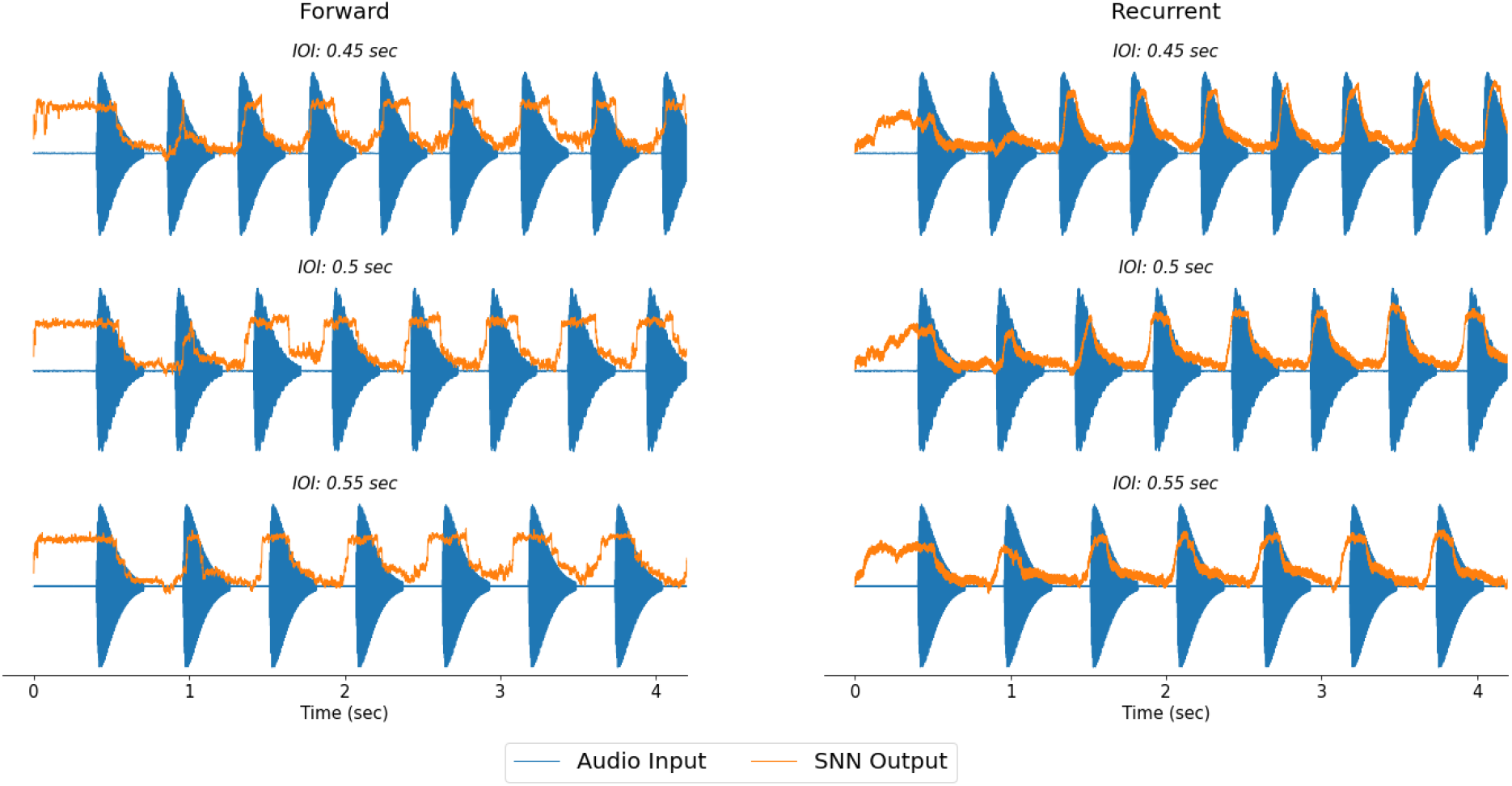
Results for Feedforward (left) and Recurrent (right) networks in the generalization task. IOI=0.45 (top row), 0.5 (mid row) and 0.55 (bottom row). Both networks were able to entrain to multiple isochronous rhythms.

As in the isochronous task, the networks exhibited oscillatory membrane-potential dynamics entrained to the beat. Figure 9 shows circular trajectories of the membrane potentials for IOIs of 0.45, 0.5, and 0.55 in the second layer. The feedforward network showed IOI-dependent changes in trajectory amplitude, whereas trajectories in the recurrent network were offset from the origin. PSTH analysis revealed similar predictive dynamics in both firing rates and membrane potentials, with exponential increases or decreases that plateaued before the predicted onsets (Supplementary Figure 2).

**Figure 9.**
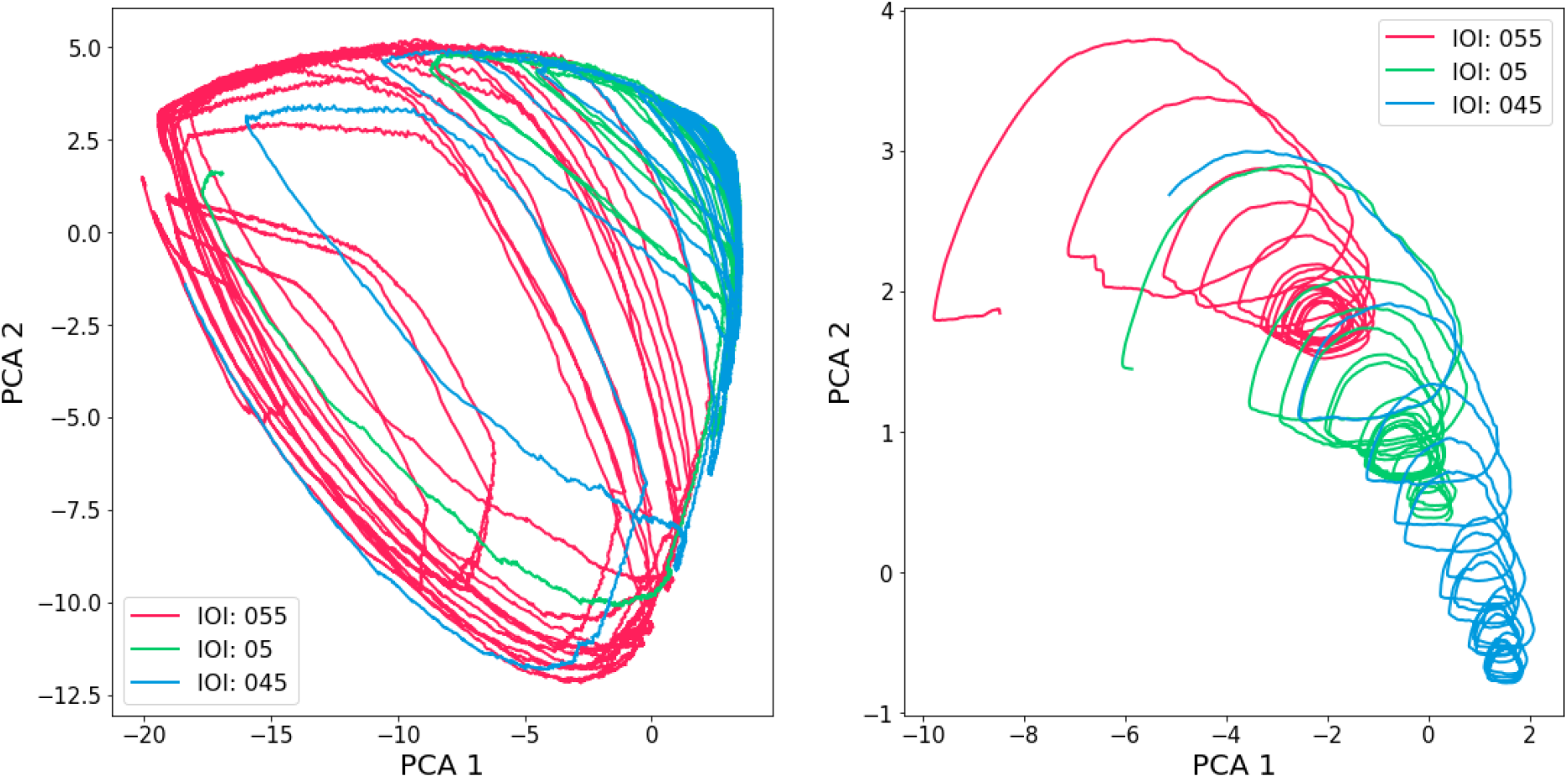
First two principal components of membrane potentials for feedforward (left) and recurrent (right) networks in the generalization task. IOIs evaluated are 0.45, 0.5, and 0.55, spanning the set of pulse intervals on which the networks were trained, and membrane potentials from the second layer are shown. These components explained over 90% of total variability in both cases. Circular trajectories are traced by these components, highlighting that the networks entrain to the input beat.

### 3.3 Mean Asynchrony

Average mean asynchrony was computed for each network that we trained, measuring how aligned the output onsets of the network were with the predicted onsets (Tables 1 and 2). Also, Figure 10 shows the asynchrony distributions for networks trained under the isochronous conditions. We observed that mean asynchrony was not uniformly distributed around zero (i.e., network output onset aligned with predicted onset), being skewed either positively (Feedforward network with IOIs of 0.4 and 0.6, Recurrent network with IOIs of 0.4 and 0.5) or negatively (Feedforward network with IOI of 0.5, Recurrent network with IOI of 0.6). Even when asynchrony was positive, it still fell below the 100 ms delay, indicating that the network output continued to anticipate the delayed onset.

**Figure 10.**
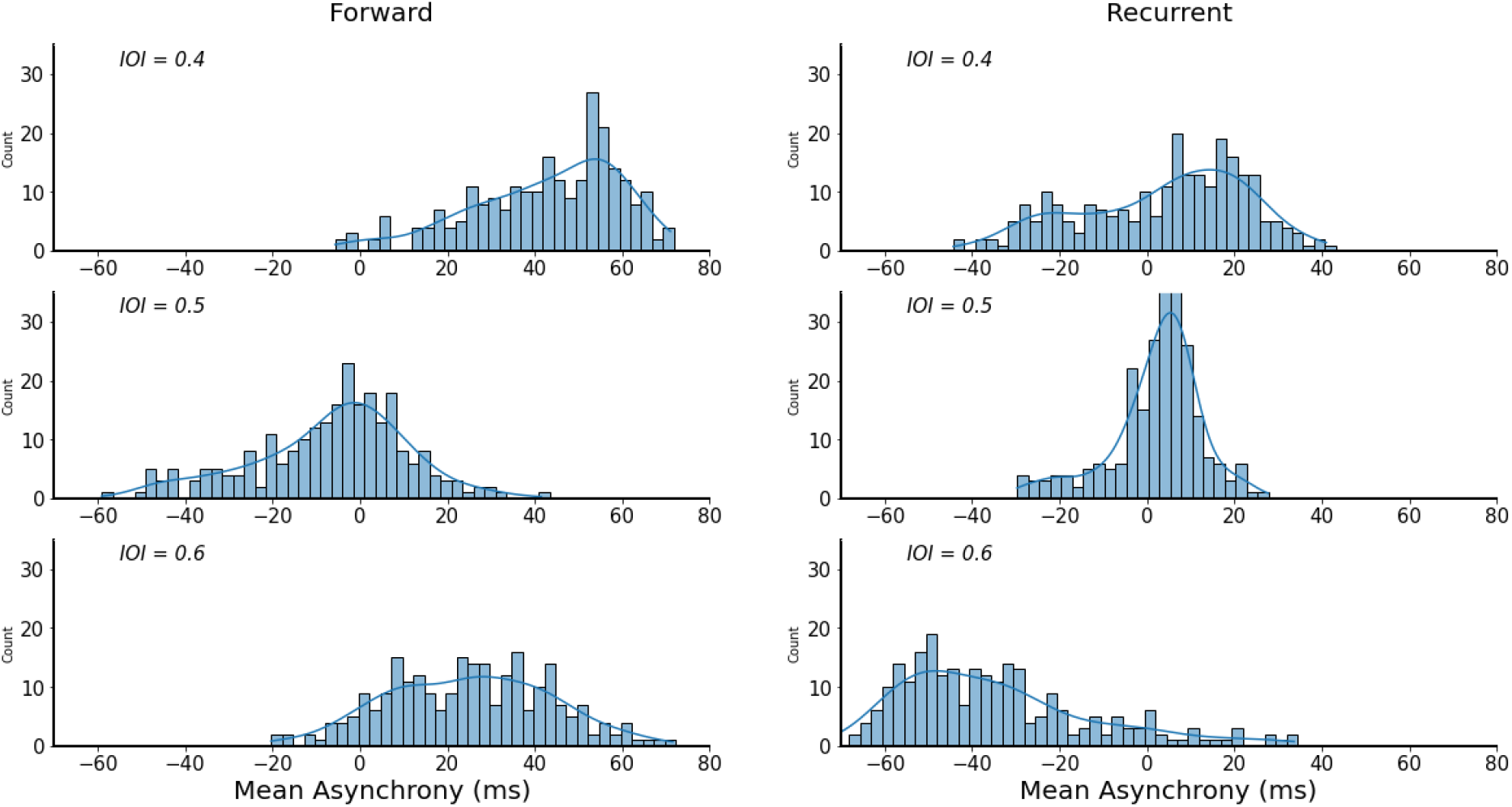
Mean asynchrony distributions of Feedforward (left) and Recurrent (right) networks for the Isochronous task. Mean asynchrony was not symmetrical. For most networks asynchrony was negative, with the exception of the feedforward network trained on IOI=0.5 s.

Further testing showed that there was a dependency of mean asynchrony on the pulse decay rate, which quantified how fast the pulse decayed after the initial onset (Fig. 1c). Figure 11 shows mean asynchrony as a function of pulse decay, and we observed that mean asynchrony depended on the pulse duration. As τ (decay rate) increases, asynchrony also tends to increase in all cases, except for the recurrent network with an IOI of 0.6. This may indicate insufficient training of this particular network, despite the high accuracy reported in Table 1. In addition, pulse frequency appeared to be uncorrelated with mean asynchrony across all cases.

**Figure 11.**
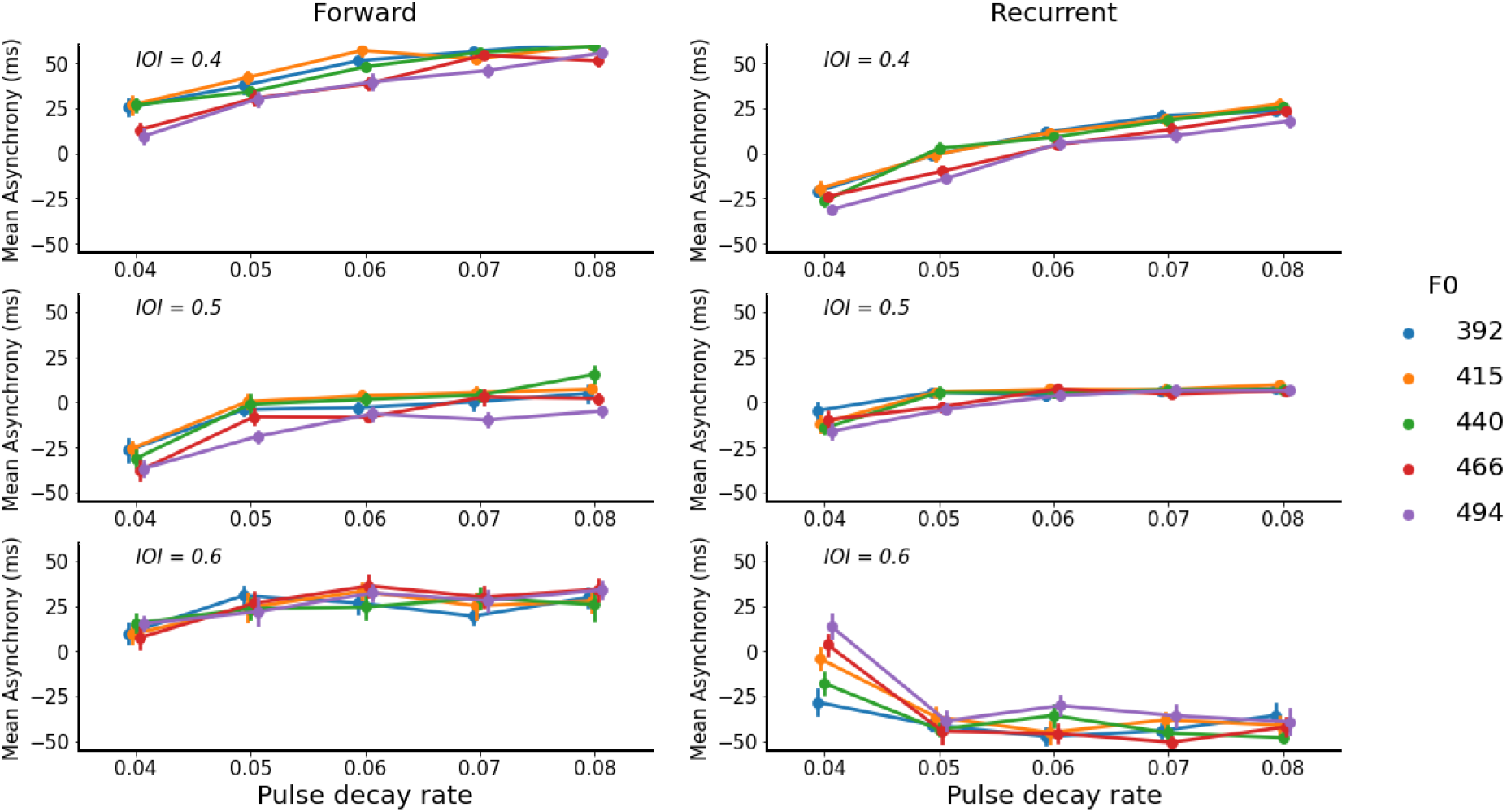
Mean asynchrony vs the pulse decay rate. A dependency between asynchrony and pulse length was observed. This dependency was different for IOI=0.6, in the network with recurrence.

When repeating the same analysis for networks that generalize across a range of IOIs (Figure 12), we observed a similar pattern. Specifically, for IOI of 0.45 seconds, mean asynchrony was positive, whereas for the other cases it was negative. Similarly, the decay rate of the pulse appeared to be positively correlated with mean asynchrony (Figure 13). Finally, as before, the fundamental frequency of the pulses did not appear to affect mean asynchrony.

**Figure 12.**
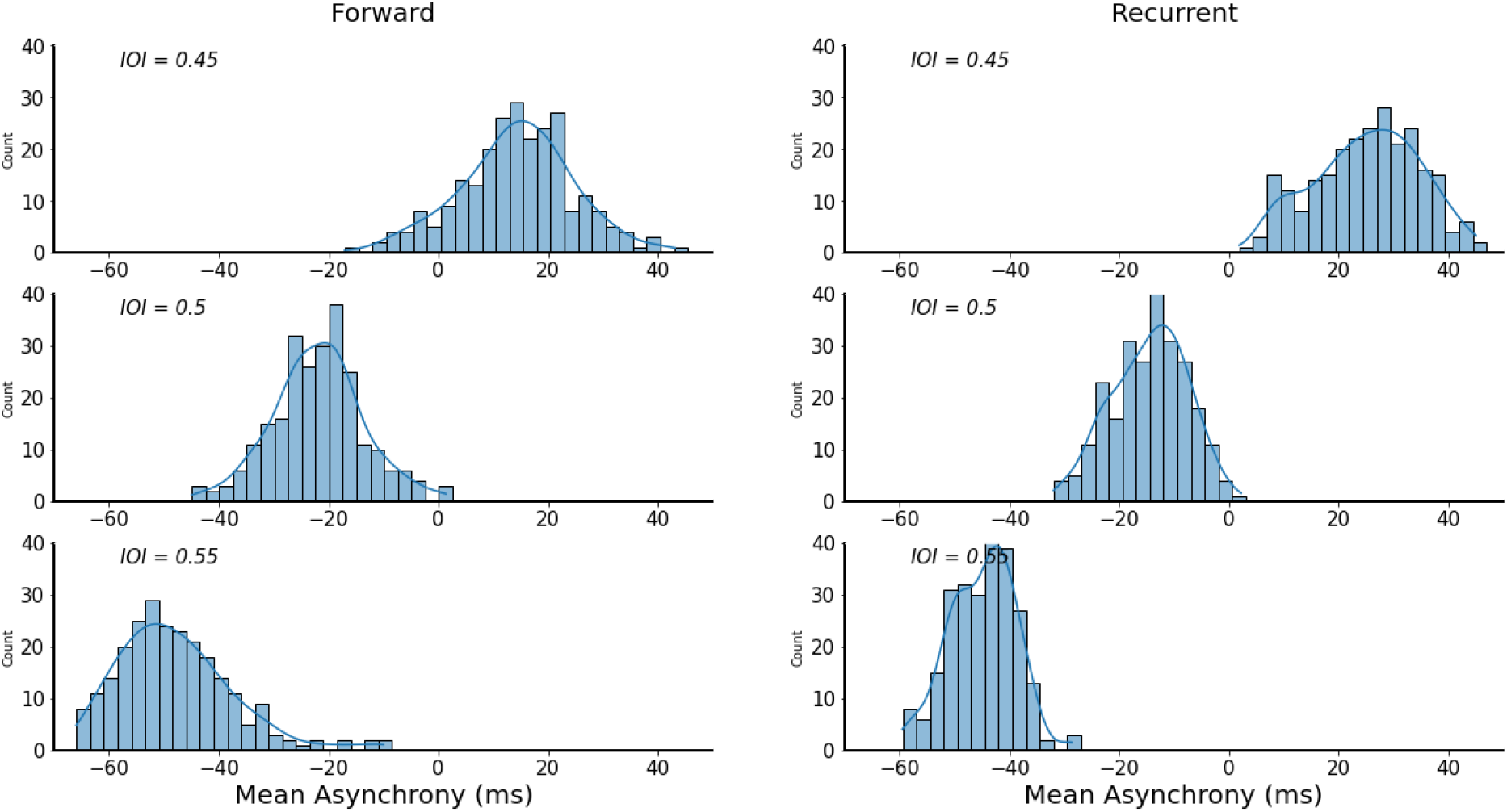
Mean asynchrony distributions of Feedforward (left) and Recurrent (right) networks trained in the generalization task. Mean asynchrony tends to increase with the IOI.

**Figure 13.**
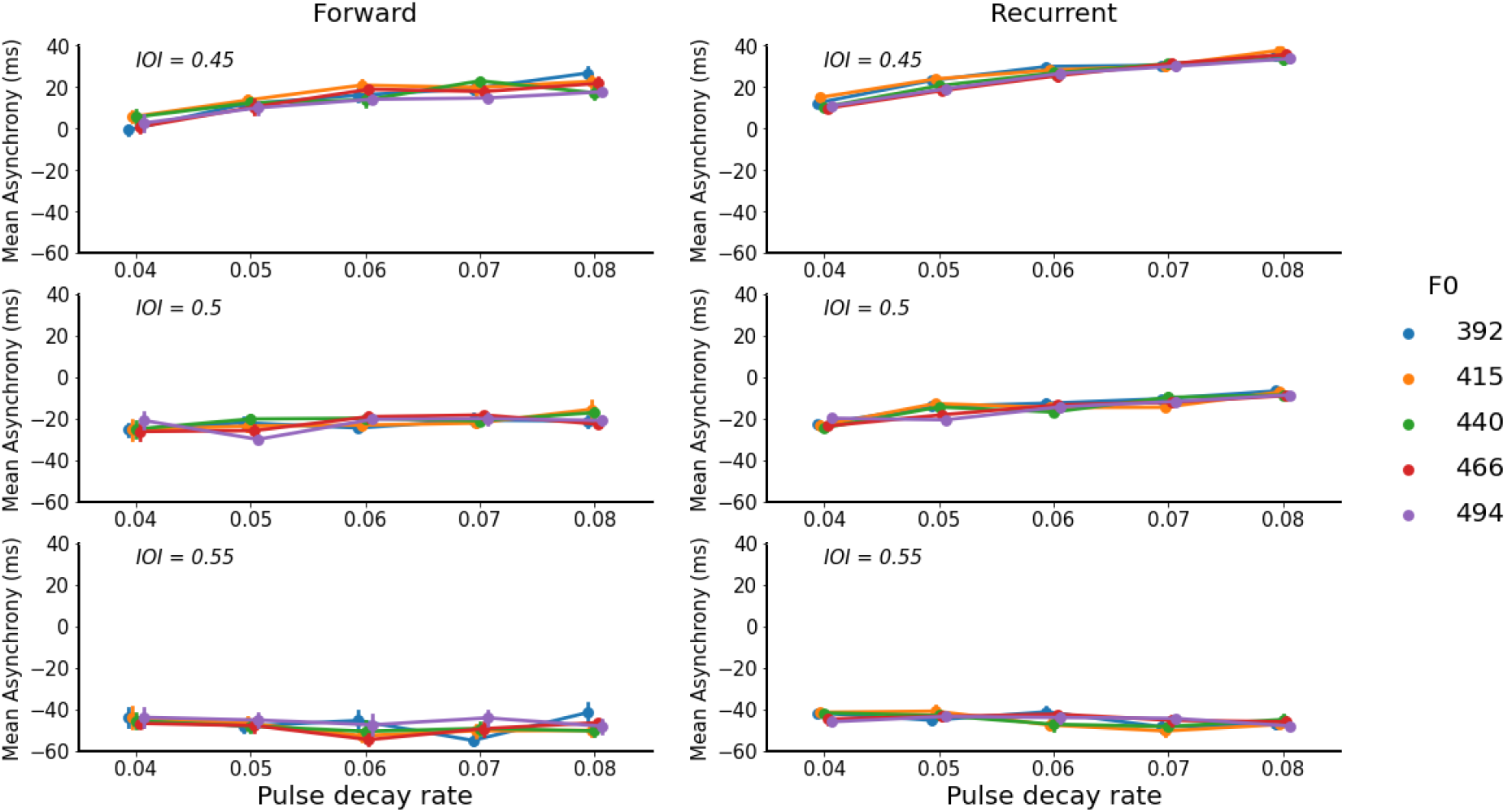
Mean asynchrony vs. the pulse decay rate of Feedforward (left) and Recurrent (right) networks similar to the isochronous case, a dependency between asynchrony and pulse length was observed. This dependency was not observed for IOI=0.55 in both cases.

Considering these results, the observed relationship between pulse decay and mean asynchrony suggests that pulse duration may contribute to triggering output onsets in the SNN. When considered alongside the gradual increase in average membrane potential around a predicted onset (Figure 7), these results support the hypothesis that different pulse durations yield different rates of membrane potential integration. This, in turn, may affect the time at which membrane potentials reach a plateau around the predicted onset, thereby influencing the precise timing of onset triggering.

## 4. Discussion

In this study, we introduced a biologically plausible spiking neural network (SNN) framework to investigate the neural basis of isochronous rhythm processing. Rather than modeling a specific region within the auditory cortex, we adopted a more abstract computational approach to investigate the mechanics and dynamics of simulated neural populations using spiking networks. We could show that a spiking neural network can entrain to incoming isochronous inputs and align to incoming event onsets with different degrees of precision. For networks trained only on specific IOIs (Figure 2), the results did not show a clear pattern of network performance and mean asynchrony (Table 1). Despite achieving an accuracy of over 90%, the asynchrony distributions varied across conditions and did not exhibit a clear relationship with performance or other network parameters. However, networks trained in the generalization case (Figure 4) showed a strong phase-tempo correlation (Table 2, Figure 11). Here, we observed that onsets lagged behind at faster tempi (IOI = 0.45) and occurred ahead of the beat at slower tempi (IOI = 0.55). Although our experiments focused on beat perception, the observed phase–tempo relationships may still relate to synchronization behavior if similar oscillatory dynamics drive activity in motor systems. In adult humans, a consistent negative onset asynchrony is observed across a wide range of tempi (Patel et al., 2005), which we did not observe here. However, the current results are more consistent with those observed in Ronan, a sea lion capable of sensorimotor synchronization (Cook et al., 2013). In an isochronous beat synchronization task (with a similar baseline tempo to ours), Ronan demonstrated beat-keeping capabilities with a strong phase-tempo correlation, where her head bobs lagged behind at faster tempi and occurred ahead of the beat at slower tempi (Cook et al., 2025). Ronan’s case is exceptional as it is one of only few documented cases of tempo flexibility outside of humans. Similarly to our networks trained on single IOIs, non-human primates can synchronize to specific isochronous stimuli but typically exhibit reactive timing (Zarco et al., 2009), although some can be trained to develop negative asynchrony (Gámez et al., 2018). However, it is worth noting that our network did not reward the emergence of negative asynchrony, and synchronization arose as a result of training under a chosen error metric.

We further showed that membrane potentials in the spiking layers entrain to the incoming beat, similarly to how neural populations entrain to musical rhythms (Nozaradan et al., 2016). At the population level, these dynamics resemble findings in non-human primates, where rhythmic tapping tasks reveal low-dimensional, oscillatory neural trajectories underlying temporal processing (Gámez et al., 2019). These results also suggest that the SNN developed coordinated population activity aligned with the temporal structure of the input. However, an important distinction lies in the role of sensory input: while primate studies reported the emergence of self-sustained dynamics in continuation tasks, the current simulations were explicitly framed within a beat perception paradigm, in which an isochronous stimulus is continuously present. Consequently, the oscillatory activity observed in the model was driven by the external input, rather than depending on fully autonomous dynamics (Zemlianova et al., 2024). Although self-sustained oscillations did not emerge under the current experimental setup, this does not rule out SNNs from generating autonomous rhythmic dynamics under different training conditions or task demands.

The current model can be compared with other rhythm models. Similar to models based on neural resonance theory (NRT; Large et al. 2015), the current results revealed that spiking layers responded to external periodic stimulation with oscillations that could phase-lock to the incoming input. Moreover, our model did not rely on an active error-correction mechanism during inference, as this only applies during training, and spiking neurons in the network were driven into an oscillatory behavior by the input. This contrasts with models that implemented an active correction mechanism to adjust internal oscillators (van der Steen and Keller, 2013) and could therefore adapt to a diverse range of inputs. One important feature of the present model is that it was based on leaky integrate-and-fire (LIF) neurons as its fundamental unit. This is similar to the model proposed by Bose et al. (2019), where spiking neurons are used as gamma counters for a beat generation model, but also relied on an active correction mechanism. Despite these differences, we hypothesized that a similar integrating (or spike-counting) mechanism should be responsible for the predictive capabilities of the model. However, the model was framed within a beat perception paradigm rather than beat generation, and in contrast to these models, this network was not designed to address rhythm generation.

We further observed an exponential increase or decrease in the membrane potentials of a subpopulation of neurons prior to predicted onsets, consistent with a leaky accumulation process used to estimate event timing. Specifically, the integration of input currents in certain neurons accumulated in the average membrane potential, producing a rapid rise until a plateau was reached. The timing of this saturation could then trigger a response in a subsequent layer, yielding the onset pulse as output. This mechanism was further explored by introducing a pulse omission (Figure 3). In this condition, the network anticipated the missing pulse, but in the absence of an input, the oscillatory response was not sustained and the network activity failed to reset. This behaviour suggests that external input may be required to initiate a new activity cycle. Without it, the neuronal response becomes effectively “stuck,” producing a mismatch between the network’s internal prediction and the actual stimulus input. Similar responses have been reported in electrophysiological studies using omission paradigms. In humans, EEG recordings have shown that the absence of an expected sound in an isochronous sequence can evoke mismatch-related responses (Prete et al., 2022). Likewise, electrophysiological recordings in animals demonstrated omission responses in auditory cortex neurons when an expected stimulus was omitted (Lao-Rodriguez et al., 2023). These findings suggest that the dynamics observed in the model capture aspects of predictive processing associated with rhythmic auditory input.

On top of the previously discussed phase–tempo correlation, we observed that the precise timing of the responses appeared to correlate with pulse length (Figures 11 and 13), suggesting that pulse duration may influence the integrating mechanism, with shorter pulses (associated with lower decay rates) producing responses that tend to anticipate the onsets. However, this relationship is not reported in the mean asynchrony literature, where pulse duration does not appear to systematically correlate with timing behaviour. A possible explanation for this discrepancy lies in the integrating mechanism described above: differences in pulse duration likely affect the effective input current and spike density, thereby modulating the rate at which activity accumulates and the point at which membrane potentials reach saturation and trigger a response. Importantly, the network was trained to minimize the MSE between the input envelope and the output, which encouraged the exploitation of continuous temporal features of the signal, including pulse duration to optimize performance. In contrast, behavioural synchronization in humans is largely driven by discrete onset detection, suggesting that different mechanisms are likely involved in biological systems. Nevertheless, the phase–tempo relationship observed here aligns more closely with the behaviour reported in Ronan (Cook et al., 2025), raising the possibility that similar integration-based dynamics may underlie her responses.

One main highlight of the current approach is that we did not explicitly model anticipatory or oscillatory behavior; instead, these features emerged naturally as the network attempted to solve the task of synchronizing to a delayed input. From theoretical work on rhythm, it was proposed that neural oscillations arise as a consequence of delays in a driven system (Stephen et al., 2008), a phenomenon we observed in the membrane potentials of our network entraining to incoming beats. This behavior was not observed in preliminary versions of the model without delays, in which the network simply decoded the input without developing any anticipatory response, as it is not needed without a delay. Recent work suggests that treating axonal delay as a learnable parameter improves network performance (Sun et al., 2023), indicating that the ability of biological networks to learn and manipulate axonal delays may provide greater flexibility for developing circuitry and dynamics that could allow networks to solve more complex rhythmic tasks. Additionally, recent work on spiking neural networks has shown that they can learn time-based information (Yu et al., 2025), further highlighting the potential of these frameworks for studying rhythmic dynamics.

Our approach has a number of limitations. Despite our efforts to construct a biologically plausible framework, we might have overlooked relevant biological phenomena. These include refractory periods in individual spiking neurons (i.e., the minimal time between two consecutive spikes) as well as constraints on firing rates. Moreover, we observed that under some conditions, membrane potentials increased indefinitely after the start of the simulation, which is unrealistic given the natural energy constraints of the nervous system. Introducing penalization terms for such behaviors during learning has been explored in spiking neural networks (Bittar and Garner, 2024). successfully reducing the occurrence of silent neurons and improving training stability. Similarly, we implemented only one-to-one neuronal recurrence, rather than all-to-all recurrence, in which the output of an entire layer is weighted and summed before being fed back into the same layer. While this approach could yield improved performance, it may also reduce the interpretability of neural population dynamics. Finally, future work should explore categorizing neurons as excitatory or inhibitory by constraining connection weights to take positive or negative values, respectively. This would allow for a more detailed investigation of inter-layer relationships and enable evaluation of the impact of specific oscillatory dynamics.

Despite the relatively strong performance of our model on the generalization task, further testing with a broader range of inter-onset intervals did not yield equally accurate results. Additionally, while the network entrained to the underlying beat, it was not able to discriminate between alternating pulses. We argue that these limitations are related, and likely arise from insufficient training data spanning a wider range of conditions, or from the absence of architectural mechanisms better suited to this task, such as all-to-all recurrence or learnable delays.

A natural direction for the future work would be to extend the present framework toward the study of sensorimotor synchronization (Repp, 2005). While the current model focused on perceptual timing and beat synchronization, SNNs provide a suitable computational framework for coupling perception and action through temporally precise spike-based signalling. In particular, the spiking output of the network could be used to drive motor unit models (Fung, 1993), in which individual spikes elicit twitch contractions that add up over time to produce a continuous force profile. By embedding the proposed SNN framework within a perception–action loop, we could investigate how anticipatory timing, entrainment, and delay-dependent oscillatory dynamics contribute to coordinated motor behavior.

## 5. Conclusions

In this work, we demonstrated that a biologically plausible spiking neural network model could reproduce certain neurobiological phenomena reported in the rhythm literature. In particular, the network exhibits anticipatory mechanisms and neural entrainment dynamics that resemble empirically observed features of beat perception. Beyond these results, the proposed computational framework provides an alternative for investigating how temporal dynamics emerge from neural interactions given specific modelling constraints, in contrast to other computational approaches where rhythm mechanisms are explicitly imposed by the model architecture.

From a computational modelling standpoint, this study highlights how recent advances in training methods for spiking neural networks can be leveraged to investigate questions traditionally addressed in neuroscience. By borrowing analysis approaches inspired by neural population dynamics, we show that it is possible to examine population trajectories across different network layers and gain insight into the mechanisms underlying emergent timing behavior. Overall, these results suggest that trained spiking neural networks can serve not only as task-performing models, but also as exploratory tools for studying neural dynamics and generating hypotheses about brain function.

## Acknowledgments

This work was funded by the FWO research project “Interactive vocal rhythms”, project number G034720N. Bart de Boer received funding from the Flemish Government under the *“Onderzoeksprogramma Artificiële Intelligentie (AI) Vlaanderen”* programme. Center for Music in the Brain (MIB) is funded by The Lundbeck Foundation (R469-2024-1573) and Købmand Herman Sallings Fond. A.R. is funded by the European Union (ERC grant TOHR 101041885). Views and opinions expressed are however those of the authors only and do not necessarily reflect those of the European Union or the European Research Council Executive Agency. Neither the European Union nor the granting authority can be held responsible for them.

## Notes

### Competing Interest Statement

The authors have declared no competing interest.

### Summary of Updates

Content moved to Supplementary Material to comply with word limits

