## Supplementary Material for "Emergent Entrainment and Predictive Dynamics in Bio-Inspired Spiking Neural Networks"

### S1. Regularization Methods

To stabilize training, we employed Threshold-Dependent Batch Normalization (tdBN) (Zheng et al., 2021), a normalization strategy designed for spiking neural networks. Unlike conventional Batch Normalization (Ioffe & Szegedy, 2015), tdBN normalizes inputs across both channel and temporal dimensions using parameters shared across timesteps, allowing variable-length inputs during training and evaluation. Alternative approaches include firing-rate regularization at the population (Zenke & Vogels, 2021) or single-neuron level (Bittar & Garner, 2024), as well as temporal extensions of Batch Normalization such as Batch Normalization Through Time (Kim & Panda, 2021) and Temporal Effective Batch Normalization (Duan et al., 2022). We selected tdBN because it supports variable-length inputs while minimizing the impact of normalization on subsequent analyses.

### S2. PCA Analysis

For each neuron, we extracted firing rates by counting spikes within a sliding window (n = 50 samples) using a Gaussian kernel. Membrane potentials were analyzed in parallel. Following methods used in finger tapping experiments (Gamez et al., 2019), we applied PCA separately to firing rates and membrane potentials at each network layer. Here, the activity of all neurons in a layer was represented by a matrix X of size [N,T], where N denotes the number of neurons and T the number of time samples. PCA yielded a transformation matrix P such that:

$$(1) \qquad Y = PX$$

where each dimension of Y explains the variance of X in descending order. We computed a specific matrix $P$ for each layer in every network, which is unique to its respective condition. For simplicity, in the isochronous task, we use an arbitrary signal with the corresponding IOI to compute the matrix $P$ to illustrate the strong periodicity of the principal components. However, in the generalization task, we required a fixed matrix $P$ across different IOIs to make a valid comparison. We simulated an evaluation signal composed of IOIs = [0.45, 0.5, 0.55], with a duration of 5 seconds for each IOI, resulting in a total duration of 15 seconds, and computed the matrix $P$ for this case. Using this $P$ ensured that the same transformation was applied to different activity patterns, and the resulting components defined trajectories that revealed the evolution of neural activity dynamics over time, allowing us to investigate how they aligned with the input.

### S3. Auditory Nerve Transduction

Auditory inputs were converted into spike trains using a phenomenological model of auditory nerve (AN) fibers (Zilany et al., 2009, 2014). Each fiber was tuned to a specific characteristic frequency (CF), producing time-varying firing rates that were converted into spike trains using a Poisson spike generator. High-spontaneous-rate fibers were simulated with CFs ranging from 125 Hz to 4 kHz and distributed logarithmically across the frequency range. The model incorporates cochlear compression, firing-rate saturation, and adaptation mechanisms (Zilany et al., 2009). A total of 32 AN fibers were simulated, providing sufficient coverage of the hearing range while maintaining computational efficiency.

Simulations were performed in *MATLAB R2022b* using a sampling rate of 100 kHz, corresponding to the minimum sampling rate supported by the model. The resulting spike trains were downsampled to 2 kHz for subsequent simulations. Downsampling was implemented by partitioning the original spike trains into non-overlapping windows of 50 samples; if a spike occurred within a window, the corresponding sample in the downsampled signal was assigned a spike. Given the sparse nature of auditory nerve firing, this procedure preserved nearly all spikes while substantially reducing computational cost.

Multiple spike-train realizations could be generated for a single auditory input due to the stochastic nature of the Poisson spike generator. Although individual realizations differed in spike timing, they preserved the same underlying firing-rate statistics. We exploited this property as a data augmentation strategy, generating multiple spike-train sets from the same auditory input.

### Supplementary Figures

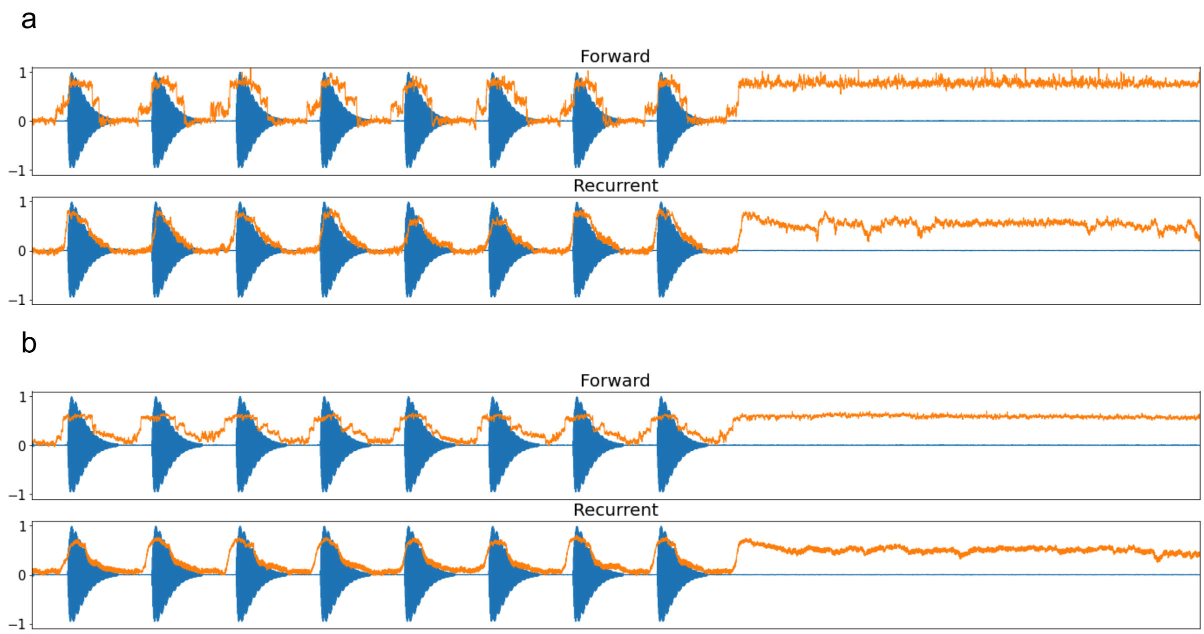


Supplementary Figure 1. Output of the model for a synchronization-continuation task. Networks are not able to sustain oscillation after the input, whether they were trained in this task or not. No oscillatory behaviour is observed on the output a) Examples for the Forward and Recurrent networks trained on IOI=0.5. b) Examples for the Forward and Recurrent networks trained on the isochronous task, for an IOI=0.5.

a

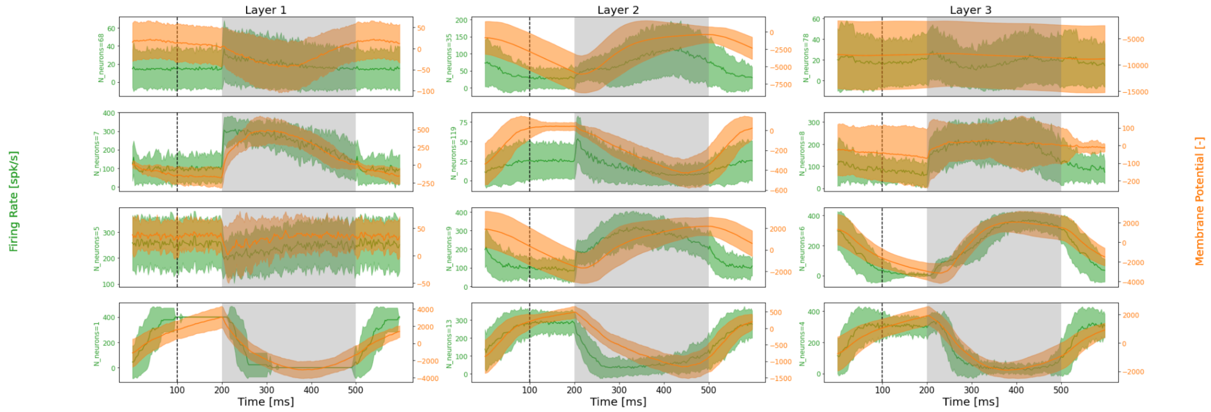


b

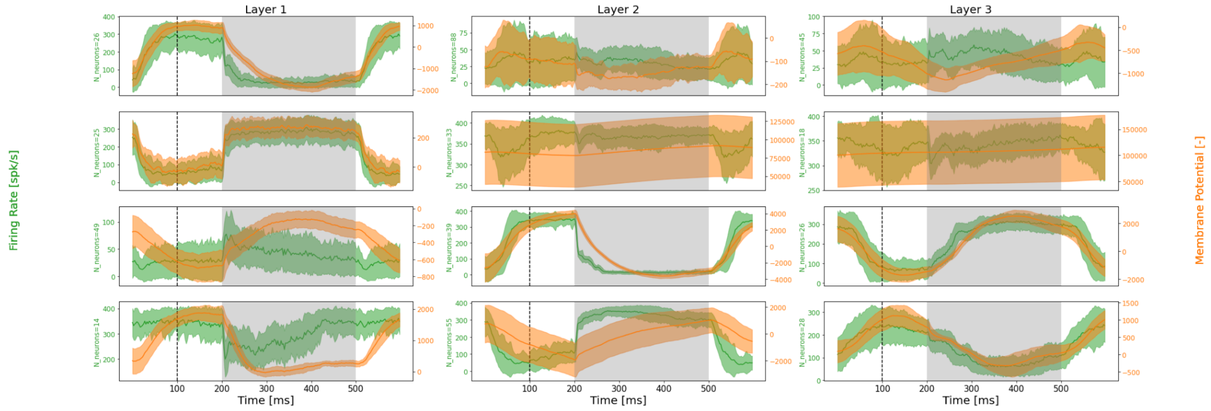


Supplementary Figure 2. PSTH of firing rates for Forward (a) and Recurrent (b) networks in the generalization task for IOI=0.5. Similar to networks trained on the Isochronous task, membrane potential and firing rates exhibit increasing or decreasing rates that pear or plateau around predicted onsets.
